# Near-complete Lokiarchaeota genomes from complex environmental samples using long and short read metagenomic analyses

**DOI:** 10.1101/2019.12.17.879148

**Authors:** Eva F. Caceres, William H. Lewis, Felix Homa, Tom Martin, Andreas Schramm, Kasper U. Kjeldsen, Thijs J. G. Ettema

## Abstract

Asgard archaea is a recently proposed superphylum currently comprised of five recognised phyla: Lokiarchaeota, Thorarchaeota, Odinarchaeota, Heimdallarchaeota and Helarchaeota. Members of this group have been identified based on culture-independent approaches with several metagenome-assembled genomes (MAGs) reconstructed to date. However, most of these genomes consist of several relatively small contigs, and, until recently, no complete Asgard archaea genome is yet available. Large scale phylogenetic analyses suggest that Asgard archaea represent the closest archaeal relatives of eukaryotes. In addition, members of this superphylum encode proteins that were originally thought to be specific to eukaryotes, including components of the trafficking machinery, cytoskeleton and endosomal sorting complexes required for transport (ESCRT). Yet, these findings have been questioned on the basis that the genome sequences that underpin them were assembled from metagenomic data, and could have been subjected to contamination and other assembly artefacts. Even though several lines of evidence indicate that the previously reported findings were not affected by these issues, having access to high-quality and preferentially fully closed Asgard archaea genomes is needed to definitively close this debate. Current long-read sequencing technologies such as Oxford Nanopore allow the generation of long reads in a high-throughput manner making them suitable for their use in metagenomics. Although the use of long reads is still limited in this field, recent analyses have shown that it is feasible to obtain complete or near-complete genomes of abundant members of mock communities and metagenomes of various level of complexity. Here, we show that long read metagenomics can be successfully applied to obtain near-complete genomes of low-abundant members of complex communities from sediment samples. We were able to reconstruct six MAGs from different Lokiarchaeota lineages that show high completeness and low fragmentation, with one of them being a near-complete genome only consisting of three contigs. Our analyses confirm that the eukaryote-like features previously associated with Lokiarchaeota are not the result of contamination or assembly artefacts, and can indeed be found in the newly reconstructed genomes.

## INTRODUCTION

Lokiarchaeota, previously referred to as Deep-Sea Archaeal group or Marine Benthic Group B, were originally detected in hydrothermal vent sites in Japan (Takai and Horikoshi, 1999) and benthic marine sediments of the Atlantic Ocean (Vetriani et al., 1999). Since then, members of this group have been found in a wide range of marine and terrestrial anaerobic/micro-aerophilic aquatic habitats, including cold seep systems, inland lakes and cave systems among others (Jørgensen et al., 2013; Sorensen and Teske, 2006). Archaea belonging to this group are metabolically active, as is suggested by the isolation of their 16S rRNA from sediments (Biddle et al., 2006) and preliminary meta-transcriptomics analyses (Cai et al., 2018). Recently, a representative of this group, ‘*Candidatus* Prometheoarchaeum syntrophicum MK-D1’ was successfully grown in co-culture showing cocci-shaped cells with long and branching protrusions (Imachi et al., 2019).

The Lokiarchaeota phylum belongs to the recently proposed Asgard superphylum together with the uncultivated phyla Thorarchaeota, Heimdallarchaeota, Odinarchaeota and Helarchaeota (Seitz et al., 2019; Seitz et al., 2016; Zaremba-Niedzwiedzka et al., 2017). Despite one complete Lokiarchaeota genome sequence is available, no complete genomes are available for any other Asgard lineages. Hence, most of our understanding of these organisms comes from metagenome-assembled genomes (MAGs) that have been reconstructed for various members of this superphylum. Phylogenomic analyses based on the reconstructed MAGs indicate that Asgard archaea affiliate with eukaryotes (Seitz et al., 2019; Spang et al., 2015; Zaremba-Niedzwiedzka et al., 2017), making this superphylum of great significance for understanding the origin of eukaryotes. Reconstructed genomes of Asgard archaea members, including Lokiarchaeota, encode numerous proteins that were previously thought to be specific to eukaryotes, so-called eukaryotic signature proteins (ESPs). These ESPs include, amongst others, several ESCRT homologs, cytoskeletal components and an expanded set of small GTPases (Klinger et al., 2016; Spang et al., 2015; Zaremba-Niedzwiedzka et al., 2017). Nevertheless, the quality and accuracy of published Asgard archaea MAGs has been challenged, including the authenticity of eukaryotic-like genes identified from these genomes, (Da Cunha et al., 2017; Garg et al., 2019). Despite several lines of evidence indicating that the position of the Asgard superphylum with respect to the eukaryotes and the presence of the so-called eukaryotic signature proteins in Asgard genomes are not the result of contamination or assembly artefacts (Spang et al., 2018), obtaining complete, high-quality Asgard archaea genomes should provide the final piece of evidence to close the debate.

Recent advances in long-read DNA sequencing technologies have made long read-based metagenomic sequencing efforts a feasible option (Hao et al., 2018; Nicholls et al., 2019). Long DNA sequencing reads improve the ability to resolve repetitive regions of sequences in *de novo* assembly, improving the contiguity of genomic assemblies. In metagenomes, repetitive sequences can be present within a single genome, as well as within several genomes that share similar regions (e.g., closely related strains, horizontally transferred genes). Therefore, if reads are long enough to span across repetitive regions, and also cover regions that are unique to a particular genome, their inclusion in metagenome assemblies can help to separate the genomic sequences of different organisms, reducing the number of chimeric contigs (Somerville et al., 2019). Furthermore, provided enough sequencing depth, metagenomic assemblies of long-read sequence datasets are expected to generate longer contigs than short-read-based assemblies, reducing potential risks of misclassifying contigs in metagenome binning efforts. Conversely, the high error rate associated with current long-read sequencing technologies creates additional challenges in the assembly process that need to be properly addressed (Nicholls et al., 2019). In particular, high error rates hinder the assembly of highly similar strains as the differences between the genomes can be indistinguishable from sequencing errors (Bertrand et al., 2019). Although several long-read assembly methods have been successfully used for single-genome *de novo* assembly (Chin et al., 2016; Koren et al., 2017) and new promising tools are being developed for metagenomic assembly (Kolmogorov et al., 2019a), how well these methods perform with real metagenomic data still remains to be properly evaluated. Yet, an increasing number of long read-based metagenomics studies demonstrate that it is possible to reconstruct complete or near-complete genomes from mock communities and natural microbial communities of varying complexity (Bertrand et al., 2019; Hao et al., 2018; Moss and Bhatt, 2018; Nicholls et al., 2019; Somerville et al., 2019).

Here we used long and short read DNA sequencing of complex microbial samples from marine sediments, to reconstruct six near-complete Lokiarchaeota genomes. Our results show that long read-based metagenomic approaches have the potential to produce near-complete genomes of low abundance lineages in complex samples, even in the presence of strain-level microdiversity. In addition, the highest quality genome we recovered here, for a species of Lokiarchaeota, provides strong evidence to that genes for ESPs identified in Asgard archaea genomes are not the result of contamination or other metagenomic artifacts, but that they are truly present in these organisms. Moreover, the long-read datasets generated in this study could also be used to validate and aid the development of metagenomics-specific methods for longread sequencing data.

## RESULTS AND DISCUSSION

### Recovery of contiguous genomes for low-abundance lineages in complex metagenomes

Various members of the Lokiarchaeota phylum have previously been found in marine sediments from Aarhus Bay (Webster et al., 2011; Zaremba-Niedzwiedzka et al., 2017). In the present study, we collected marine sediments using a Rumohr core from the same Aarhus Bay site sampled previously (Zaremba-Niedzwiedzka et al., 2017). Sediment from this newly obtained core was sub-sampled at five-centimeter intervals, beginning five centimeters below the sea floor (cmbsf), until the lowest depth of 65 cmbsf. The relative abundance of 16S rRNA amplicon reads belonging to the Lokiarchaeota phylum ranged from 1.1% to 7.6% depending on the sampling depth. Within all of the 5 cm interval sediment sub-samples, a total of 21 operational taxonomic units (OTUs) classified as Lokiarchaeota were identified, with relative abundances for a single OTU of up to 5.3% (Fig. 1a; Suppl. Table 1). Two samples (from 20 and 25 cm below the sediment-water interface, referred to as C20 and C25 henceforth), which showed the highest relative abundance for one of the Lokiarchaeota OTUs, were used to generate DNA sequence datasets using both long and short read technologies. To provide sufficient quantities of environmental DNA for these sequencing methods, multiple DNA extractions from C20 and C25 were performed and pooled together. Illumina sequencing was used to produce high-quality reads of short length, yielding ∼80 Gbp of usable data for C20 and ∼85 Gbp for C25 (Suppl. Table 2). Additionally, 4 runs of Oxford Nanopore sequencing (1 Promethion run for C20 and 1 Minion and 2 Promethion runs for C25) were performed resulting in ∼47 Gbp of read data for C20 and ∼61 Gbp of read data for C25 (post filtering), with median read lengths ranging between 2992 bp and 4557 bp, depending on the run (Suppl. Table 2). To verify the presence of putative Lokiarchaeota lineages in C20 and C25, an initial metagenomic assembly using the short Illumina reads was performed and proteins present in contigs containing at least 5 out of 15 ribosomal proteins (RP15) (Zaremba-Niedzwiedzka et al., 2017) were used to were used to carry out a phylogenetic analysis. Despite C20 and C25 containing the same Lokiarchaeota lineages according to the 16S rRNA amplicon analysis, the RP15 phylogeny showed 8 and 5 contigs containing Lokiarchaeota ribosomal proteins in C20 and C25, respectively (Fig. 1b), suggesting that the abundance of some Lokiarchaeota lineages in C25 was too low to assemble the ribosomal operon with the data generated. The subsequent assemblies were performed using reads generated from the C25 sample as they had the highest Lokiarchaeota abundance and produced longer Lokiarchaeota contigs in initial long-read assemblies, and reads from the C20 sample were only used to aid binning.

**Table 1.** Summary statistics for the hybrid assemblies and for the SR-L04 MAG generated from Illumina reads. Statistics for the complete genome belonging to the MK-D1 strain are included as a reference.

|  | **L15** | **L04** | **L11** | **L02** | **L05** | **L03** | **SR-L04** | **MK-D1** |
| --- | --- | --- | --- | --- | --- | --- | --- | --- |
| **MAG size (Mbp)** | 4.45 | 4.07 | 4.13 | 3.39 | 3.77 | 3.50 | 3.13 | 4.43 |
| **Number of contigs** | 9 | 3 | 6 | 36 | 21 | 48 | 398 | 1 |
| **Average contig length (bp)** | 494473 | 1355997 | 688058 | 94276 | 179490 | 72905 | 7854 | 4427796 |
| **Median contig length (bp)** | 401335 | 1729042 | 693839 | 65251 | 115125 | 48503 | 4991 | 4427796 |
| **N50 (bp)** | 446705 | 2146130 | 693839 | 116878 | 314162 | 121159 | 12106 | 4427796 |
| **Largest contig (bp)** | 1496208 | 2146130 | 1173416 | 403150 | 609122 | 351171 | 68064 | 4427796 |
| **Shortest contig (bp)** | 150292 | 192821 | 182219 | 15710 | 16462 | 16407 | 1509 | 4427796 |
| **GC (%)** | 31.45 | 32.68 | 32.63 | 31.87 | 29.89 | 30.71 | 32.69 | 31.17 |
| **Number of Ns** | 52 | 100 | 100 | 457 | 600 | 216 | 0 | 0 |
| **N (%)** | 0 | 0 | 0 | 0.01 | 0.02 | 0.01 | 0 | 0 |
| **CDSs** | 4202 | 3826 | 3934 | 3583 | 3807 | 3717 | 2886 | 3880 |
| **rRNAs** | 3 | 3 | 3 | 3 | 3 | 3 | 1 (partial 16S) | 3 |
| **Completeness (default) (%)** | 88.79 | 87.85 | 87.38 | 80.73 | 88.55 | 76.32 | 70.09 | 91.74 |
| **Contamination (default) (%)** | 3.74 | 2.8 | 6.39 | 0.47 | 6.07 | 19.73 | 1.87 | 6.07 |
| **Strain Heterogeneity (default) (%)** | 0 | 0 | 44.44 | 0 | 12.5 | 36.36 | 0 | 9.09 |
| **Completeness (reduced) (%)** | 95.92 | 94.9 | 94.39 | 87.13 | 94.64 | 82.31 | 75.51 | 96.09 |
| **Contamination (reduced) (%)** | 2.04 | 2.04 | 5.44 | 0 | 5.1 | 21.04 | 1.02 | 5.1 |
| **Strain Heterogeneity (reduced) (%)** | 0 | 0 | 66.67 | 0 | 20 | 38.1 | 0 | 16.67 |

**Figure 1.**
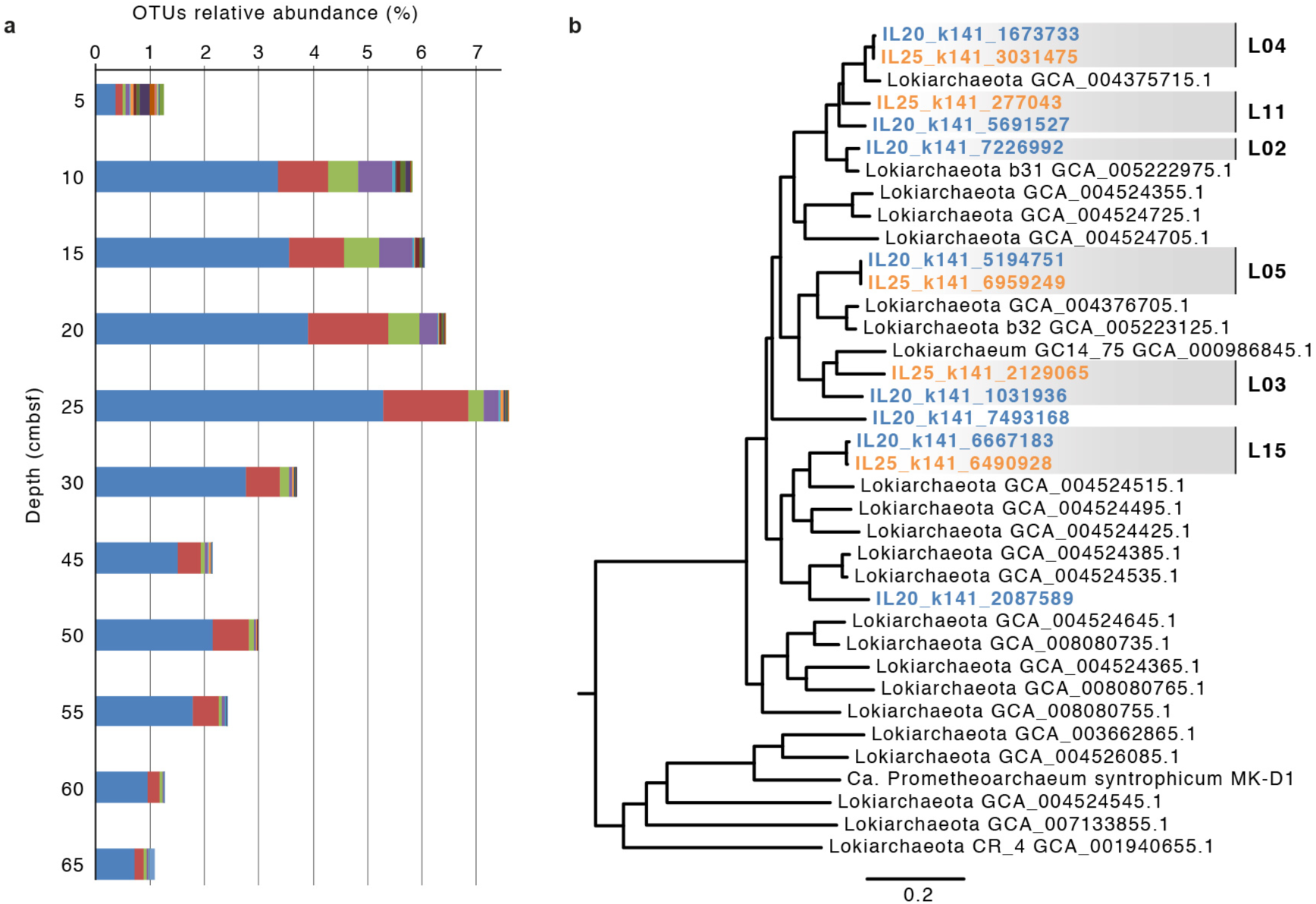
Exploration of Lokiarchaeota abundance and diversity. **a,** 16S rRNA amplicon-based assessment of Lokiarchaeota diversity and abundance across different depths from the M5 sampling site in Aarhus Bay. Colors represent different Lokiarchaeota operational taxonomic units. Relative abundance values are estimated based on the total (prokaryotic and eukaryotic) diversity. Measurements in centimeters correspond to depths below the water-sediment interface at which the sediment core was sub-sampled and used for DNA extractions. **b,** Phylogenetic diversity of Lokiarchaeota in samples C20 (blue) and C25 (orange) based on short reads-derived contigs that contain at least 5 out of 15 ribosomal proteins (see Methods). Main clades representing putative lineages for which MAGs were reconstructed are highlighted in grey.

As the aim of this study was to generate complete, or near-complete, genomes of specific taxa and not to assemble the whole community, we followed a strategy that aimed to reduce the amount of data generated, whilst also enriching our datasets for reads sequenced from Asgard archaea genomes. One benefit of reducing the number of sequencing reads is that it allowed us to test various long-read sequence assemblers without reaching computational limitations. In fact, data reduction is a common step in various long-read assemblers, in which certain fraction of the longest reads is selected prior to assembly. Even though the selection of the longest sequencing reads could help to achieve better results for projects aiming to assemble individual genomes, data reduction based on read length could also negatively impact metagenomics assemblies of datasets that contain low-abundance community members. For this reason, we decided to avoid length-based filtering in the present study.

The overall strategy we followed consisted of four main steps: database generation, read recruitment, assembly and binning (Fig. 2a). First, a custom database of Asgard archaea genomes was created by gathering available MAGs, and by generating new MAGs from the C25 dataset. In a first iteration, MAGs were obtained by assembling and binning all short and long sequencing reads independently (Fig. 2b). Megahit was used to assemble the short-read Illumina datasets. For the long-read Nanopore data, only one assembly could successfully be generated using all available long-reads with the computational resources available using miniasm (Li, 2016). A second assembly, which only included reads longer than 4000 bp, was generated using Marvel (github.com/schloi/MARVEL) producing highly contiguous assemblies at the expense of creating several evident chimeric contigs between some lineages. All assemblies were binned separately using Metabat2 (Kang et al.) and coverage information from samples C20 and C25. MAGs belonging to the Asgard superphylum were subsequently identified and included in the custom Asgard archaea genome database (Fig. 2b). Other variations in the database construction were also performed to attempt to maximize the number of recruited read sequences from Asgard archaea genomes. Second, reads were classified by homology search and kmer classifications against the local database described above (Fig. 2c; see Methods for details). Next, the recruited reads were combined into contigs using several assemblers and subsequently binned. For each lineage that could be clearly identified, we manually selected the most contiguous MAGs out of all the assemblies generated, avoiding those with evident chimeras likely created by the co-assembly of closely related lineages. Chimeric co-assemblies were especially common between lineages L04 and L11, and between L03 and L05. The performance of the different assemblers in the presence of closely related lineages varied considerably and, in several cases, parameter settings markedly affected the results. The selected contigs were used in a final iteration to separate reads originated from the different Lokiarchaeota lineages and assemble them individually using the hybrid assembler Masurca (Zimin et al., 2017), as this assembler produced better results in terms of overall consensus sequence and contiguity (Fig. 1d). This strategy resulted in the generation of 6 Lokiarchaeota MAGs with bin sizes ranging between 3.4 and 4.5 Mb (Fig. 1b; Table 1). To reduce the number of frameshift and errors present in the final set of MAGs, contigs were further polished using short reads. All 16S sequences identified in the MAGs belonged to the same OTU generated from the amplicon data over comparable regions (percentage of identity higher than 97%), which highlights the limitations of amplicon classifications to estimate the abundance of closely related organisms. Yet, local assembly errors in the 16S region cannot be excluded.

**Figure 2.**
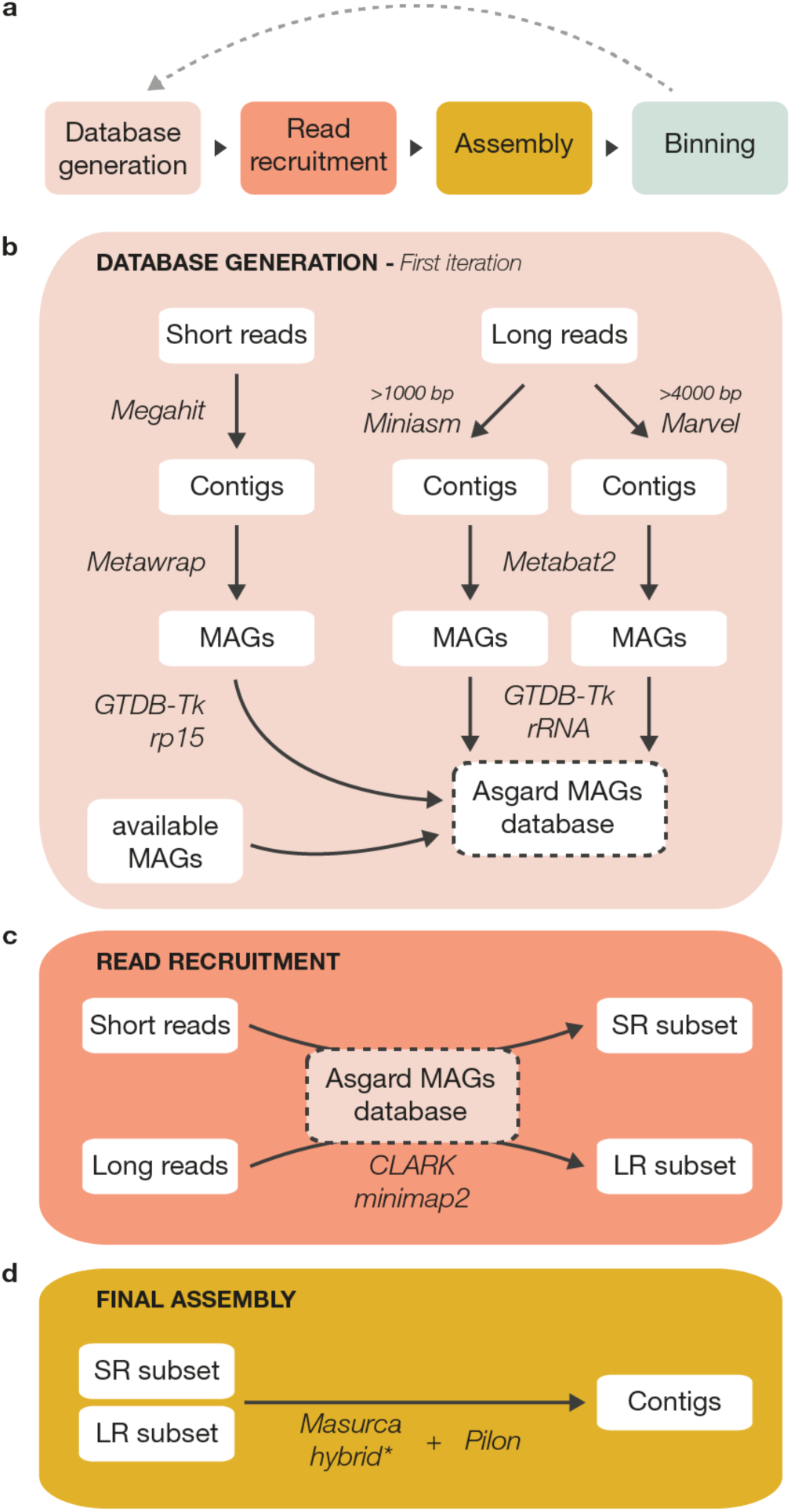
Assembly workflow. **a,** A schematic overview of the assembly approach. **b,** The steps followed to generate a custom database of Asgard archaea (draft-)genomes. The database was iteratively modified to incorporate subsequently binned genome data, following successive iterations of read recruitment and genome assembly. **c,** Outline of the read recruitment approach. **d,** Tools used to generate the final hybrid assembly. Asterik indicates that the step was performed individually for each lineage.

### A near-complete Lokiarchaeota genome

In order to base our analyses on the most robust data available, we first identified which of the six assembled Lokiarchaeota MAGs had the best quality. This was determined as the MAG corresponding to Lokiarchaeota L04, since this genome had the highest coverage and was assembled in the least number of contigs (three), with a combined total length of 4.1 Mb. Hybrid MAGs from L11 and L15 were also highly complete but slightly more fragmented and with higher levels of contamination. For L04, the initial estimated completeness and contamination values of this genome according to CheckM (Parks et al., 2015) were 89% and 3% respectively (Table 1). However, since some marker genes are suggested to be absent in Lokiarchaeota genomes (Narrowe et al., 2018), we decided to create a Lokiarchaeota-specific set of marker genes by excluding those absent or duplicated in the three most complete MAGs reconstructed in the present study (L04, L11 and L15), in order to obtain more reliable estimations of completeness and contamination for this phyla (Suppl. Fig. 1). Diphthamide biosynthesis proteins were among the missing marker genes (Suppl. Table 3), in agreement with previous studies that suggest they are missing from Lokiarchaeota genomes (Narrowe et al., 2018). After correction for these missing genes, the completeness and contamination values estimated using the Lokiarchaeota marker gene set were 95% and 2% for L04, respectively (Table 1). Additionally, independent estimations of completeness and contamination values for the complete genome of the Lokiarchaeota strain ‘*Ca.* Prometheoarchaeum syntrophicum MK-D1’ based on conserved archaeal marker genes were suboptimal (91% and 6% respectively), further emphasizing the need of a cautious interpretation of these values.

To determine whether there were any major assembly errors and chimeric contigs in the L04 hybrid genome, we first aligned both the short and long reads against the genome and inspected the changes in coverage depth. Although long reads are less prone to changes in coverage than short reads, there were still several regions of coverage variation across the genome. In metagenomic datasets, changes in coverage could be explained by unspecific read alignments (e.g., from other organisms or repetitive/conserved regions), genetic variation within populations (i.e., genomic regions only present in a subpopulation), or actual mis-assemblies (i.e., regions incorrectly reconstructed that do not represent any real genomic sequence). To minimize non-specific and partial read alignments, we created a subset of long read alignments (filtered-alignment hereafter), that include only those alignments spanning at least 85% of the read length and with an identity value compared to the hybrid genome of at least 85%. Given the high Nanopore error rates (∼14-20%) (Weirather et al., 2017), such high identity cut-off might result in the removal of true read-alignments but the stringency of this subset can be helpful to identify regions incorrectly reconstructed.

We searched within the genome for bases not covered by any long read from the unfiltered alignment, as such cases would indicate clear problems within the assembly. Unsurprisingly, we found few regions with zero coverage close to the ends of contigs, suggesting that these regions might be erroneous (Suppl. Table 6). Mis-assemblies located at the end of contigs are a common issue in *de novo* genome assembly, not necessarily indicating a problem with the overall assembly. In particular, the end of one of the contigs (C1S) seemed to be especially problematic showing a drop in read coverage that overlaps with a region containing several proteins with bacterial best hits in sequence similarity searches (Fig. 3). This pattern could either be caused by the co-assembly of two different organisms (i.e., a chimeric region incorrectly reconstructed from genomic information of two different organisms), or by biological differences at subpopulation-level, such as a recent horizontal transfer, present in some members of the population but missing in others. However, given that some bases in the assembly sequence are not covered by any reads, the former scenario seems to be more likely. Additionally, we identified two other main regions, R1 and R2 of ∼30 and ∼20 Kbp respectively, which contained several bases with no or low coverage in the filtered long-read alignment (Fig. 3). Examination of the reads aligned to regions R1 and R2 could not determine the cause of this coverage pattern. Future generating of additional and longer read data might help to resolve the sequence of these regions more accurately (Suppl. Fig. 2a and b).

**Figure 3.**
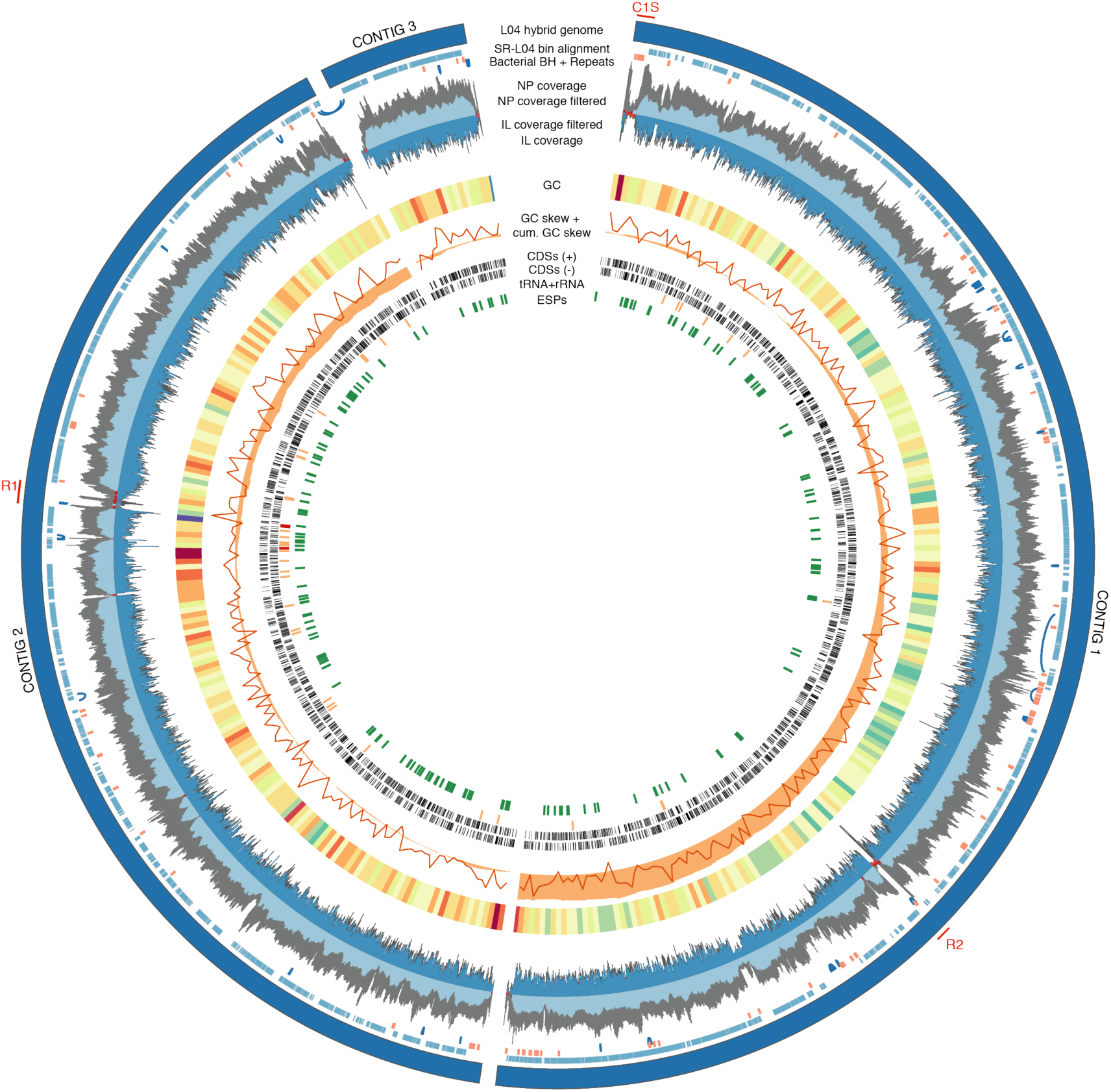
Representation of the Lokiarchaeota L04 genome. From outside to inside: 1. Hybrid assembly (dark blue); 2. Alignment of the SR-L04 bin; 3. Proteins with bacterial best hits (orange) and repeats identified by mummer self-hits (blue). Crossing repeats are omitted. 4. Nanopore coverage before (grey) and after (light blue) filtering. 5. Illumina coverage before (dark grey) and after (blue) filtering; 6. GC content heatmap in which dark red represent AT rich areas and dark blue/purple represents GC rich areas; 7. GC-skew (orange line) and cumulative GC-skew (orange histogram); 8 and 9. CDS in the positive and negative strand, respectively; 10. tRNAs (orange) and rRNAs (red); 11. Eukaryotic signature proteins (green).

None of the assembly tools tested in this study were able to generate a complete genome for any Lokiarchaeota lineage. Different tools often resulted in incompatible genome reconstructions suggesting problems in the current algorithms and highlighting that manual inspection is still necessary to obtain accurate genome assemblies. The development and validation of specialized tools for *de novo* metagenomic assembly of long-reads will likely improve current assemblies. Likewise, future improvements in DNA extraction methods that allow the recovery of less fragmented DNA from challenging environmental samples (such as marine sediments), will result into longer sequencing reads that will definitely help to produce more contiguous assemblies and resolve long repeats.

Future analyses that focus on improving the hybrid assemblies generated in this study by using information from read-alignments, inspection of repetitive sequences and integration of alternative genome reconstructions generated with different assemblers might lead to close and more accurate genome reconstructions of these lineages. Additionally, targeted PCR followed by Sanger sequencing can be used to confirm the genomic sequence of various regions, or to attempt to close the genome whenever it is not possible to do it bioinformatically with the current data available.

### A MAG generated from short reads is highly fragmented but accurate

For the L04 strain, we investigated how a non-curated MAG that was generated by binning the contigs from the assembly of short Illumina reads without any further bin-refinement step (SR-L04 hereafter), compared to the L04 hybrid genome. To do this, the contigs previously assembled from Illumina reads were binned using two different tools – metabat2 (Kang et al., 2019) and maxbin2 (Wu et al., 2016) – and the results were combined using binning_refiner from metawrap (Song and Thomas, 2017; Uritskiy et al., 2018). The total size of SR-L04 was 3.13 Mb and it was comprised of 398 contigs with an average length of 7854 bp (Table 1). The high level of fragmentation observed in the assembly of this genome is probably caused by the presence of closely related-lineages and microvariation in the sample. Yet, the estimated completeness was still relatively high with values of 70% and 76% according to CheckM based on the Archaea and Lokiarchaeota marker gene sets, respectively (Table 1). A similar value (75%) was obtained from aligning the SR-L04 to the higher quality and more complete L04 hybrid assembly (Suppl. Fig. 3). Consistent with this, we found that 75% of the CDSs predicted in the hybrid assembly genome were also predicted in SR-L04. Unsurprisingly, the missing fraction of the genome included regions that are often difficult to accurately assemble, such as rRNA genes and repeats-containing regions (Fig. 3). The only rRNA gene that was identified from SR-L04 was a partial 16S rRNA gene, while the hybrid assembly genome harbored a complete set of 5S, 16S and 23S rRNA genes. Interestingly, the estimated contamination for SR-L04 was very low (1.87% and 1.02% according to CheckM Archaeal and Lokiarchaeota set, respectively), even though no further manual inspection or bin refinement was performed. A similar contamination value (1.2%) was also determined, based on the percentage of bases in SR-L04 that could not be aligned to the L04 hybrid genome. In total, nine SR-L04 contigs could not be aligned to the hybrid genome. These contigs had an average length of 2022 bp and were all shorter than 3000 bp. However, since the genome reconstructed from the hybrid assembly is not entirely complete, it is possible that some or all of the nine unaligned SR-L04 contigs are actually part of the genome of this organism.

Altogether, we observed that, in spite of the fragmented nature of SR-L04, the quality of this reconstructed genome is medium-high (Bowers et al., 2017). In addition, the binning strategy used here, which exclude subsequent binning-refinement steps, seems to be relatively conservative, sacrificing completeness in favor of stringency and low contamination values. Although these results cannot be generalized to all MAGs in the metagenome, the current binning procedure was able to produce an accurate bin for a species of Lokiarchaeota from short read data alone in this particular case. However, lineages with less clear composition or abundance patterns could still cause problems in the binning step. For example, composite MAGs that included several of the other Lokiarchaeota lineages that had a lower abundance were observed (data not shown). Our results show that it is possible to generate highly accurate (albeit incomplete) MAGs even when no further bin-refinement step is performed. Hence, MAGs should not automatically be branded as artefactual and contaminated because of their metagenomic origin. The quality of MAGs can vary for different metagenomic datasets, taxa and binning approaches used, and therefore, we highlight the need for manual inspection and bin refinement to ensure their accuracy.

### Eukaryotic signature proteins in Lokiarchaeota genomes are not the result of chimeras and contamination

As reported for other Lokiarchaeota genomes previously (Spang et al., 2015; Zaremba-Niedzwiedzka et al., 2017), the Lokiarchaeota L04 hybrid assembly reconstructed in the present study is enriched in genes encoding ESPs. These included genes encoding homologues of cytoskeleton proteins, such as actin, profilin and gelsolin, as well as multiple small GTPases and components of the oligosaccharyltransferase (OST) protein complex (Suppl. Table 4). As was also shown to be the case for other Lokiarchaeota genomes (Spang et al., 2015; Zaremba-Niedzwiedzka et al., 2017), we found homologues of genes for the endosomal sorting complex required for transport (ESCRT) components, with most genes encoded in a gene cluster. As is the case for *Lokiarchaeum* sp. GC14-75, one of the ESCRT-III components is encoded in a different region of the genome that, in the L04 hybrid genome, is located next to a gene containing a steadiness box domain characteristic of the Vps23/TSG101 subunit of the eukaryotic ESCRT-I. Similarly, components of the ubiquitin protein modifier system were identified and also found to be encoded in a gene neighborhood as previously reported for other Lokiarchaeota genomes (Zaremba-Niedzwiedzka et al., 2017) (Suppl. Table 4).

The putative ESPs were grouped into seven categories: “Cytoskeleton”, “Ubiquitin”, “ESCRT”, “Trafficking machinery”, “GTPases” and “OST”. For each category, we could identify a number of ESPs (Suppl. Table 5) in the L04 hybrid genome similar to those reported for the Lokiarchaeote CR-4 genome (175 and 157, respectively) and the ‘*Ca*. Prometheoarchaeum syntrophicum MK-D1’ complete genome (131). The number of ESPs reported for the *Lokiarchaeum* sp. GC14-75 genome was slightly higher (211), which is expected as this genome is known to be redundant, containing several closely related strains in the same MAG (Spang et al., 2015). The identified ESPs were widely dispersed throughout the L04 hybrid genome (Fig. 3), as was previously shown to be the case for *Lokiarchaeum* sp. GC14-75 (Spang et al., 2015). Additionally, we investigated the read coverage for these regions and saw no signs of coverage anomalies with the exception of a putative ubiquitin-conjugating enzyme found at the contig end in the C1S region, which was flagged as problematic on the basis of low read coverage and presence of multiple proteins with bacterial best hits in homology searches (Fig. 3).

## CONCLUSIONS

In this study we show that is possible to reconstruct near-complete genomes for low-abundant taxa in complex microbial communities using a combination of long and short read sequencing technologies. The availability of long sequencing reads was essential to obtain long contigs that were otherwise highly fragmented in the assemblies generated with short Illumina sequencing reads alone. Using this data, we were able to reconstruct a near-complete Lokiarchaeota MAG, consisting of only three contigs, together with five additional MAGs that were slightly more fragmented. Our analyses show that the ESPs previously reported in Lokiarchaeota are present in the highest-quality assembly recovered for this clade from the present study. Our findings therefore confirm that the presence of ESPs in Lokiarchaeota is not the result of binning and assembly artefacts.

## MATERIALS AND METHODS

### Data availability

The Lokiarchaeota MAGs obtained from hybrid assemblies using both short- and long-read metagenomics (Illumina and Nanopore) datasets are available via this link: https://figshare.com/articles/figshare_tar_gz/11378847.

### Sediment sampling and DNA extraction

A sediment sample was collected from sampling station M5 (56° 06′ 12′′ N, 10° 27′ 28.2′′ E) at Aarhus Bay, Denmark, using a Rumohr core. 2 ml sediment subsamples were then collected from the core at 5 cm vertical intervals, beginning at 5 cm below the sediment-water interface. DNA was extracted from each sediment subsample using the DNeasy PowerLyzer PowerSoil kit (QIAGEN) in accordance with the manufacturer’s protocol.

### Universal 16S rRNA gene amplicon sequencing and OTUs generation

‘Universal’ primer pairs A519F (5’-CAGCMGCCGCGGTAA-3’) and U1391R (5’-ACGGGCGGTGWGTRC-3’) were used to amplify 16S rRNA genes using reaction conditions specified previously (Spang et al., 2015). Barcoded amplicon sequencing libraries were constructed as described previously (Spang et al., 2015) prior to sequencing with an Illumina MiSeq instrument (2×300 bp). Reads were processed to remove primer sequences and bases at the 3’ end with a Phred quality score < 10 using cutadapt v1.10 (Martin, 2011), leaving reads of at least 100 bases. Forward and reverse reads were de-replicated and clustered into centroid OTUs independently using VSEARCH v. 1.11.1 (--derep_fulllength; threshold=97%) (Rognes et al., 2016). UCHIME (Edgar et al., 2011) with the SILVA132_SSUref_tax:99 database (Quast et al., 2013) was used to remove chimeric reads. The remaining reads were taxonomically classified using the LCAClassifier from CREST-3.0 (Lanzén et al., 2012) with silva132 as the reference database (Quast et al., 2013).

### Short-read library preparation and sequencing

Libraries were created by the SciLifeLab SNP&SEQ Technology Platform using the ThruPLEX DNA-seq library preparation kit (Rubicon Genomics). Illumina paired-reads of length 150 bp were generated on a Illumina NovaSeq instrument.

### Long-read library preparation and sequencing

Originally, a single MinION sequencing run (NP25m1) was performed to produce long reads for C25. However, the amount of data obtained was insufficient to generate an adequate depth of coverage, required to produce long contigs, for any Lokiarchaeota lineage. Consequently, two additional Promethion runs were generated, one from C25 (NP25p1) and another for C20 (NP20p1). Unfortunately, the PromethION run for the C25 data failed and produced a limited amount of data and, thus, an additional Promethion run for that sample (NP25p2) was required. The same DNA extraction described above for C25 and C20 was used to generate the sequencing library required for MinION (NP25m1) and Promethion (NP20p1) sequencing. For the following long-read sequencing runs (NP25p1 and NP25p2) the DNA was extracted using the DNeasy Powersoil kit (QIAGEN) as it has a slightly gentler lysis step. In order to obtain 1 μg of DNA required for long-read sequencing several DNA extractions were pooled together. High molecular weight (HMW) DNA extraction method using the MagAttract HMW DNA kit, (QIAGEN) was attempted but unsuccessful as particles in the sediment bound to the magnetic beads used to isolate HMW DNA. All long-read sequencing libraries were carried out from 1 μg of DNA using the SQK-LSK109 kit (Oxford Nanopore Technologies, Oxford, UK) according to the manufacturer’s protocol. Nanopore sequencing was performed using FLO-MIN106 and FLO-PRO002 flowcells for MinION and PromethION respectively. MinION reads were basecalled using the Albacore v2.3.3 with the r94_450bps_linear.cfg configuration. Promethion basecalling was done real-time in MinKNOW, using Guppy v1.8.5.

### Read preprocessing

Adapters and low-quality bases present in Illumina reads were trimmed using Trimmomatic v0.33 with the following parameters: PE ILLUMINACLIP:NexteraPE-PE.fa:2:30:10 SLIDINGWINDOW:4:15 MINLEN:100 (Bolger et al., 2014). Nanopore reads shorter than 1000 bp and with a mean accuracy lower than 80% were filtered using FiltLong v0.2.0 -- min_lenght 1000 --min_mean_q 80 (github.com/rrwick/Filtlong). Adapters were removed if present using Porechop v0.2.3_seqan2.1.1 (github.com/rrwick/Porechop).

### Short-read sequencing data assembly and binning

Illumina reads were assembled using megahit v1.1.3 with default parameters (Li et al., 2016). Contigs longer than 1000 bp were binned using metawrap v1.2 (Uritskiy et al., 2018) including Illumina reads from samples C20 and C25 with the options --metabat2 --maxbin2 --universal --run-checkm and refined with the bin_refinement module and the options -c 50 -x 10. MAGs belonging to the Asgard superphylum (Asgard MAGs) were identified with GTDB-Tk v0.2.2 (github.com/Ecogenomics/GTDBTk). Furthermore, contigs containing a subset of ribosomal proteins (RP) were classified by aligning and concatenating RPs as explained in (Zaremba-Niedzwiedzka et al., 2017) and inferring a phylogeny with FastTree v2.1.10 (Price et al., 2010). Contigs encoding ribosomal proteins branching within the Asgard archaea superphylum were used to identify additional Asgard MAGs.

### Long-read sequencing data assembly and binning

Assemblies using the full set of Nanopore reads were performed using Minimap2 v2.16-r922 and Miniasm v0.3-r179 (Li, 2016, 2018) (including reads >= 1000 bp) and Marvel (Git commit 7885338) (github.com/schloi/MARVEL) (including reads >= 4000 bp). For each assembly, binning was performed with metabat2 v2.12.1 (Kang et al.) using short and long reads from samples C20 and C25. MAGs belonging to the Asgard superphylum were identified with GTDB-Tk v0.2.2 (github.com/Ecogenomics/GTDBTk). Additionally, 16S and 23S rRNA genes were identified using Barrnap v0.9 (github.com/tseemann/barrnap) and phylogenies were inferred using FastTree v2.1.10 (Price et al., 2010). MAGs containing rRNA genes branching within Asgard were further selected.

### Read recruitment and re-assembly

A database of genomes and MAGs belonging to the Asgard superphylum was generated by combining the Asgard MAGs previously identified with other available Asgard genomes. Variations of the database were performed by including or excluding newly binned MAGs from various read subsets. This database was used to classify Illumina and Nanopore reads with CLARK v1.2.3 (Ounit et al., 2015) and Minimap2 v2.16-r922 (Li, 2018). The following assemblers were tested on the read subsets: Canu v1.8 (Koren et al., 2017), Flye v2.4.2 (Kolmogorov et al., 2019b), Ra v0.2.1 (github.com/lbcb-sci/ra), Wtdbg2 v.2.4 (Ruan and Li, 2019), Masurca (Zimin et al., 2017), Unicycler v0.4.7 (Wick et al., 2017), Marvel (github.com/schloi/MARVEL), OPERA-MS (Bertrand et al.) and HINGE (Kamath et al., 2017). Assemblies were manually inspected to select the longest contigs for each Lokiarchaeota strain avoiding those with clear chimeras. The selected contigs were used to identify the short and long reads originated from each individual strain in a new run of read-recruitment. For each strain, the recruited reads were assembled with the hybrid assembler Masurca (Zimin et al., 2017). The final contigs were combined and polished altogether using 3 iterations of pilon v.1.22, prior short-reads alignment using bowtie2 v3.4.3 (Langmead and Salzberg, 2012) with the following parameters: --local --very-sensitive-local -I 0 -x 2000.

### Estimation of genome completeness and contamination

Genome completeness and redundancy were estimated with CheckM v1.0.5 (Parks et al., 2015) using the archaeal set of marker genes. Additionally, a Lokiarchaeota-specific subset of marker genes was derived by excluding duplicated and absent genes (PF04010, PF01866, TIGR00289, TIGR03679, TIGR00336, TIGR00522, TIGR03677, TIGR03685, PF01287, PF00679, PF00867, PF00958) in the three most contiguous genomes (L04, L11 and L15).

### Estimation of genome coverage

Short reads were aligned to MAGs using bowtie2 v3.4.3 (Langmead and Salzberg, 2012). More stringent read-mapping was derived by filtering proper read pairs that had less than 3 mismatches to the reference using hts_nim_tools bam-filter v0.0.1 (github.com/brentp/hts-nim-tools). Long reads were aligned to MAGs using Minimap2 v2.16-r922 (Li, 2018), excluding secondary alignments using samtools v1.9 (Li et al., 2009). More stringent read alignment was derived by selecting reads with a query coverage equal or greater than 85% of the query length and an identity value of at least 85%. Visualization of the reads alignments was done using IGV v2.4.17 (Thorvaldsdóttir et al., 2013) and Ribbon (Nattestad et al., 2016 n.d.). Per-base coverage was calculated using bedtools genomecov v2.27.1 (Quinlan and Hall, 2010). Average per-window depth was computed using mosdepth 0.2.5 (Pedersen and Quinlan, 2018) with a window size of 1000 bp and visualized using circos v0.69.6 (Krzywinski et al., 2009).

### MAGs comparison

SR-L04 contigs were aligned against L04 contigs using bwa mem v0.7.17 with the option -x intractg (Li et al., 2009) to calculate the number of aligned bases in both the query and the subject. Circular visualization of alignments between SR-L04 and L04 was performed using jupiter minBundleSize=500 (github.com/JustinChu/JupiterPlot). Dotplot and global alignment were created using mummerplot v.3.5 prior alignment of the contigs using nucmer v.1 with the --nosimplify option (Kurtz et al., 2004).

### Identification of previously reported ESPs

Homology searches of the predicted CDSs against all proteins from *Lokiarchaeum* sp. GC14-75 and Lokiarchaeote CR-4 were performed with diamond v0.9.24 (Buchfink et al., 2015). Proteins aligned to ESPs sequences previously reported in (Spang et al., 2015) and (Zaremba-Niedzwiedzka et al., 2017) were selected as putative ESPs if the e-value < 1e-6 and a percentage of the query coverage of at least 70%. The depth of coverage across ESPs was calculated from the read alignments described above.

---

**Supplementary Figure 1.**
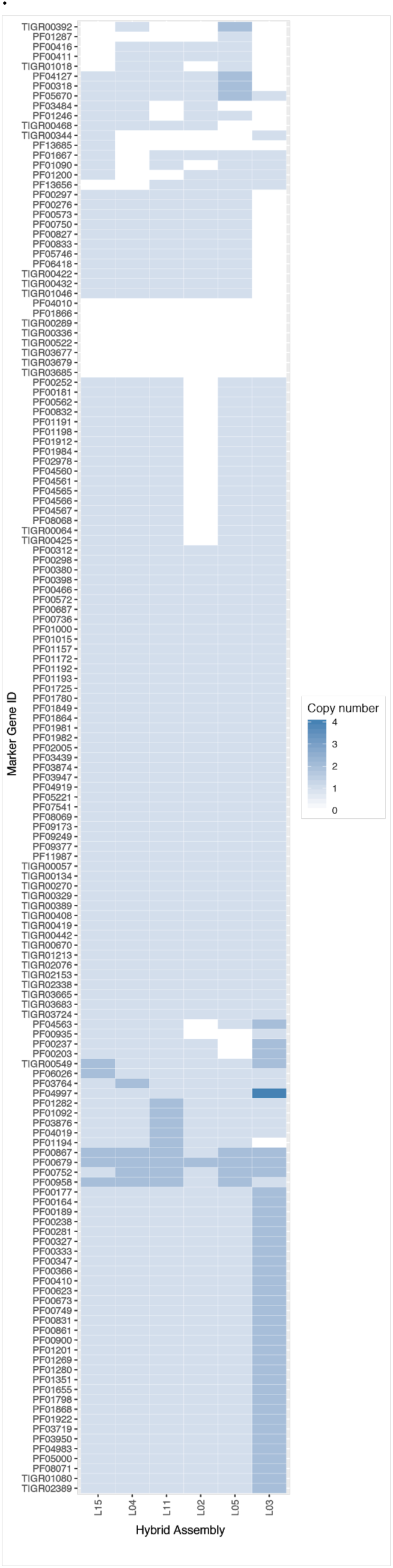
Presence of archaeal marker genes used by CheckM in the Lokiarchaeota hybrid MAGs.

**Supplementary Figure 2.**
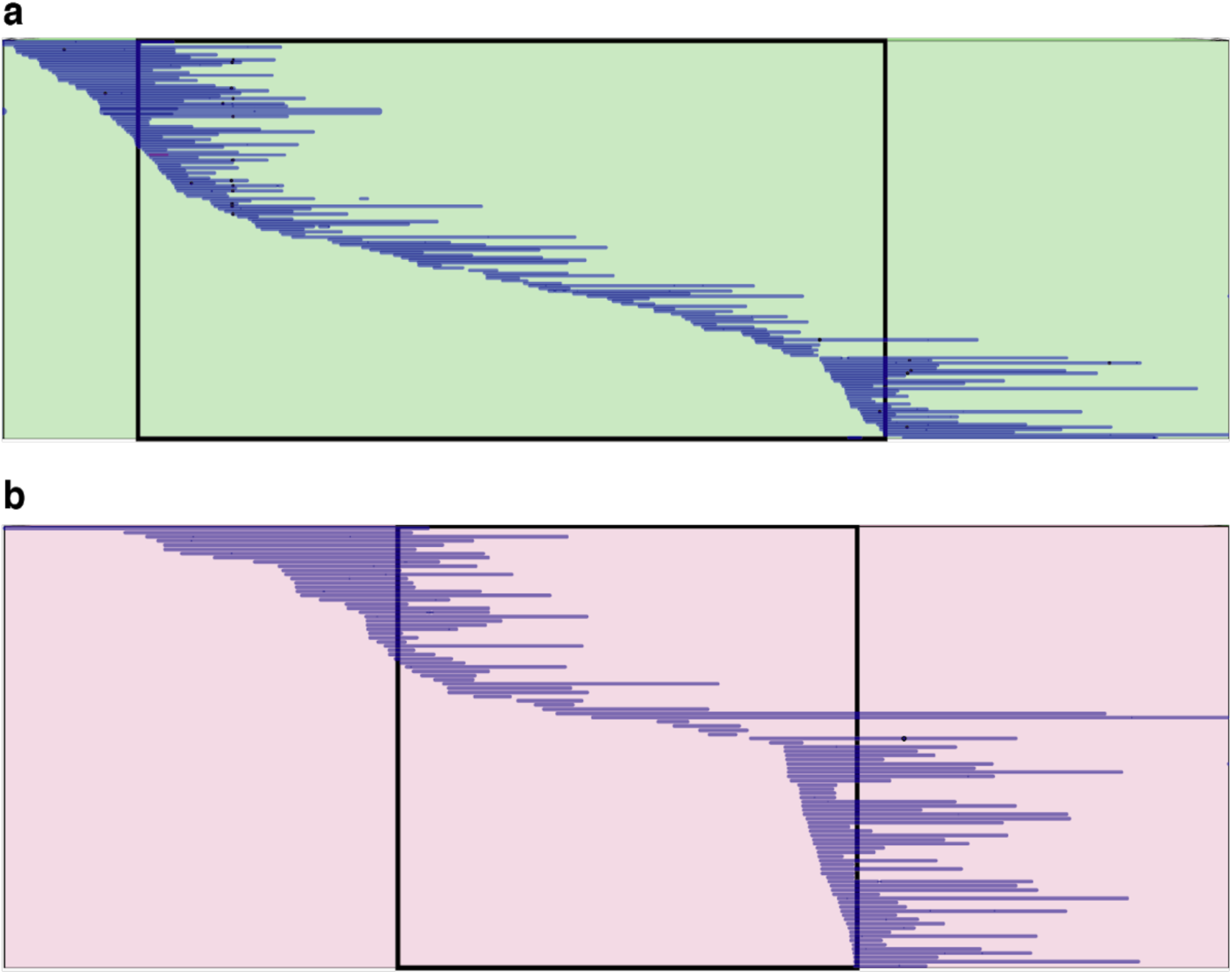
Filtered long-reads alignment across regions with potential misassemblies, R1 (a) and R2 (b), visualized with Ribbon (Nattestad et al., 2016). Blue lines represent reads alignments and black dots correspond to insertions.

**Supplementary Figure 3.**
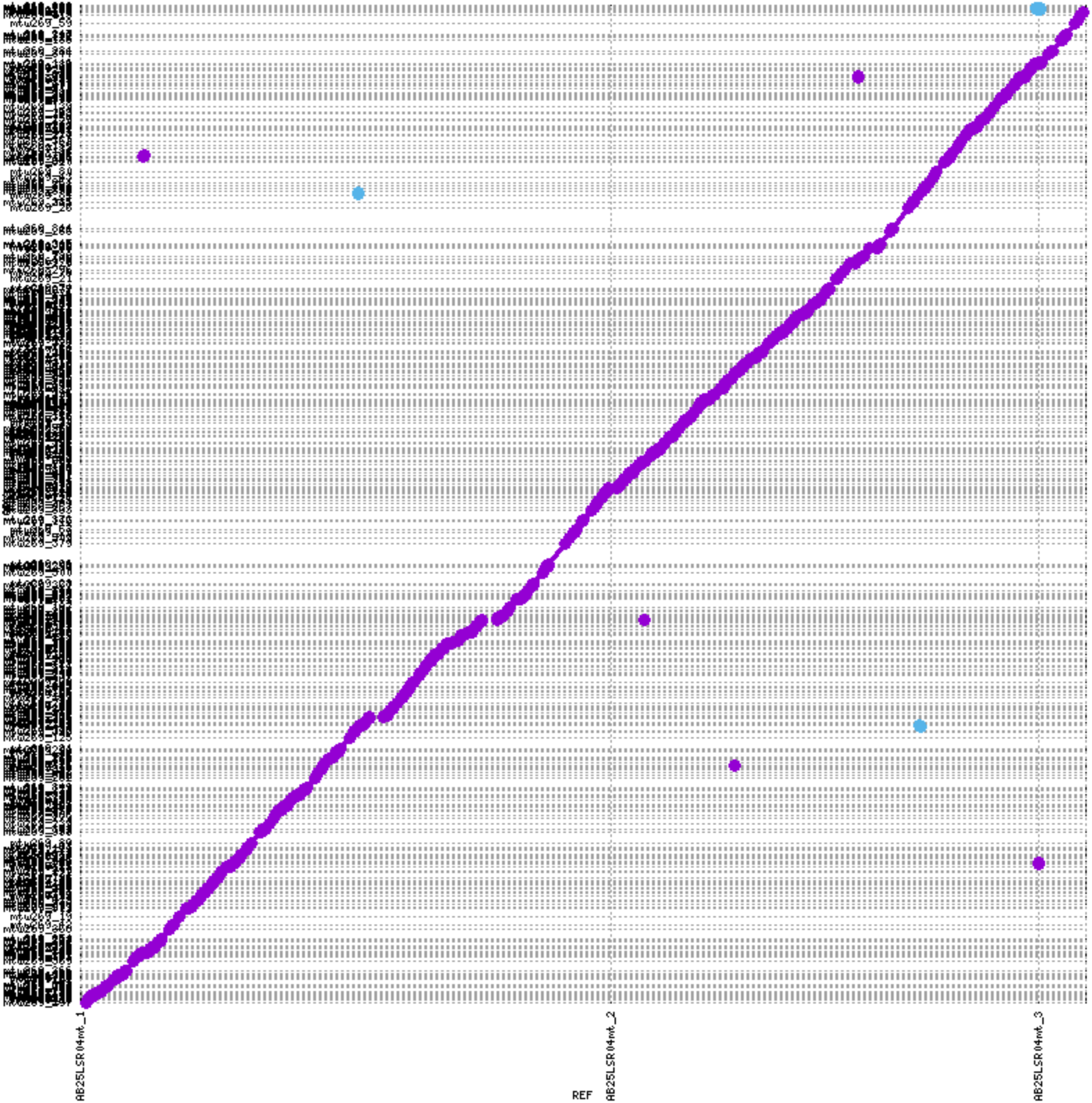
A DNA sequence alignment of the L04 hybrid genome and SR-L04, represented by a dotplot diagram. Dots show alignments in the same (purple) and reverse direction (blue). Axis represent contig sequences in the L04 hybrid genome (x axis) and SR-L04 (y axis).

**Supplementary Table 1.** Relative abundance of Lokiarchaeota OTUs at the sampling site M5 (Aarhus Bay) across various depths.

|  | 5 cm | 10 cm | 15 cm | 20 cm | 25 cm | 30 cm | 45 cm | 50 cm | 55 cm | 60 cm | 65 cm |
| --- | --- | --- | --- | --- | --- | --- | --- | --- | --- | --- | --- |
| OTU_99_9471 | 0.37<br>0 | 3.34<br>3 | 3.54<br>8 | 3.90<br>4 | 5.28<br>1 | 2.76<br>9 | 1.50<br>1 | 2.15<br>0 | 1.77<br>6 | 0.95<br>0 | 0.71<br>1 |
| OTU_1820_142<br>1 | 0.12<br>9 | 0.92<br>2 | 1.02<br>6 | 1.48<br>0 | 1.57<br>0 | 0.61<br>1 | 0.42<br>3 | 0.66<br>4 | 0.47<br>9 | 0.21<br>8 | 0.17<br>5 |
| OTU_3830_248 | 0.04<br>2 | 0.56<br>4 | 0.63<br>9 | 0.57<br>8 | 0.29<br>3 | 0.18<br>0 | 0.08<br>1 | 0.09<br>3 | 0.06<br>6 | 0.05<br>5 | 0.05<br>7 |
| OTU_3293_247 | 0.08<br>7 | 0.62<br>1 | 0.61<br>7 | 0.32<br>0 | 0.27<br>3 | 0.04<br>1 | 0.06<br>2 | 0.03<br>4 | 0.05<br>8 | 0.01<br>8 | 0.02<br>4 |
| OTU_5707_10 | 0.01<br>8 | 0.06<br>1 | 0.03<br>8 | 0.02<br>1 | 0.04<br>0 | 0.01<br>3 | 0.01<br>9 | 0.00<br>9 | 0.00<br>5 | 0.00<br>0 | 0.01<br>5 |
| OTU_1561_36 | 0.04<br>8 | 0.00<br>5 | 0.00<br>7 | 0.01<br>7 | 0.04<br>0 | 0.03<br>8 | 0.02<br>2 | 0.01<br>4 | 0.01<br>2 | 0.01<br>0 | 0.01<br>0 |
| OTU_2738_15 | 0.00<br>0 | 0.01<br>0 | 0.01<br>0 | 0.01<br>0 | 0.03<br>3 | 0.01<br>3 | 0.01<br>9 | 0.00<br>4 | 0.00<br>5 | 0.00<br>5 | 0.00<br>6 |
| OTU_2169_45 | 0.05<br>5 | 0.06<br>9 | 0.06<br>8 | 0.03<br>4 | 0.01<br>9 | 0.00<br>7 | 0.00<br>2 | 0.00<br>1 | 0.00<br>1 | 0.00<br>0 | 0.00<br>0 |
| OTU_1683_37 | 0.05<br>3 | 0.09<br>2 | 0.04<br>9 | 0.04<br>1 | 0.01<br>0 | 0.00<br>8 | 0.00<br>5 | 0.00<br>0 | 0.00<br>0 | 0.00<br>0 | 0.00<br>1 |
| OTU_937_61 | 0.18<br>5 | 0.09<br>5 | 0.02<br>4 | 0.00<br>7 | 0.01<br>0 | 0.00<br>5 | 0.00<br>0 | 0.00<br>0 | 0.00<br>0 | 0.00<br>0 | 0.00<br>0 |
| OTU_6595_2 | 0.00<br>0 | 0.00<br>8 | 0.00<br>1 | 0.00<br>3 | 0.00<br>8 | 0.00<br>0 | 0.00<br>0 | 0.00<br>0 | 0.00<br>0 | 0.00<br>0 | 0.00<br>0 |
| OTU_7090_2 | 0.09<br>8 | 0.01<br>0 | 0.00<br>1 | 0.00<br>3 | 0.00<br>2 | 0.00<br>0 | 0.00<br>0 | 0.00<br>0 | 0.00<br>0 | 0.00<br>0 | 0.00<br>0 |
| OTU_1844_22 | 0.00<br>0 | 0.00<br>0 | 0.00<br>0 | 0.00<br>0 | 0.00<br>0 | 0.00<br>0 | 0.00<br>5 | 0.00<br>0 | 0.02<br>8 | 0.00<br>5 | 0.08<br>8 |
| OTU_6288_2 | 0.02<br>6 | 0.00<br>0 | 0.00<br>0 | 0.00<br>0 | 0.00<br>0 | 0.00<br>0 | 0.00<br>0 | 0.00<br>0 | 0.00<br>0 | 0.00<br>0 | 0.00<br>0 |
| OTU_5857_2 | 0.04<br>0 | 0.00<br>0 | 0.00<br>0 | 0.00<br>0 | 0.00<br>0 | 0.00<br>0 | 0.00<br>0 | 0.00<br>0 | 0.00<br>0 | 0.00<br>0 | 0.00<br>0 |
| OTU_6523_2 | 0.00<br>5 | 0.00<br>0 | 0.00<br>1 | 0.00<br>0 | 0.00<br>0 | 0.00<br>0 | 0.00<br>0 | 0.00<br>0 | 0.00<br>0 | 0.00<br>0 | 0.00<br>0 |
| OTU_5521_3 | 0.00<br>0 | 0.00<br>0 | 0.00<br>0 | 0.00<br>0 | 0.00<br>0 | 0.00<br>0 | 0.00<br>0 | 0.00<br>0 | 0.00<br>0 | 0.00<br>0 | 0.00<br>0 |
| OTU_5469_3 | 0.00<br>0 | 0.00<br>0 | 0.00<br>0 | 0.00<br>0 | 0.00<br>0 | 0.00<br>0 | 0.00<br>0 | 0.00<br>0 | 0.00<br>0 | 0.00<br>0 | 0.00<br>0 |
| OTU_5367_5 | 0.02<br>6 | 0.00<br>0 | 0.00<br>1 | 0.00<br>0 | 0.00<br>0 | 0.00<br>0 | 0.00<br>0 | 0.00<br>0 | 0.00<br>0 | 0.00<br>0 | 0.00<br>0 |
| OTU_7499_2 | 0.00<br>0 | 0.00<br>0 | 0.00<br>0 | 0.00<br>0 | 0.00<br>0 | 0.00<br>0 | 0.00<br>0 | 0.00<br>0 | 0.00<br>0 | 0.00<br>0 | 0.00<br>0 |
| OTU_5978_2 | 0.05<br>8 | 0.00<br>5 | 0.00<br>0 | 0.00<br>0 | 0.00<br>0 | 0.00<br>0 | 0.00<br>0 | 0.00<br>0 | 0.00<br>0 | 0.00<br>0 | 0.00<br>0 |
| SUM | 1.24<br>1 | 5.80<br>6 | 6.03<br>3 | 6.41<br>6 | 7.57<br>8 | 3.68<br>6 | 2.13<br>8 | 2.96<br>9 | 2.43<br>1 | 1.26<br>1 | 1.08<br>7 |

**Supplementary Table 2.** Summary statistics for the generated sequencing data after filtering.

| Sample | Sequencing instrument | Read type | Filtered reads |  |  |  |  |
| --- | --- | --- | --- | --- | --- | --- | --- |
|  |  |  | Yield (Gb) | Reads (M) | Median quality | Median read length (bp) | Longest read (Kb) |
| C25 | NovaSeq (Illumina) | 2x150 | 84.93 | 2 x 284.11 | - | - | - |
| C25 | MinION (Nanopore) | long | 7.98 | 1.5 | 9.1 | 4557 | 71.41 |
| C25 | PromethION (Nanopore) | long | 13.3 | 2.94 | 8 | 2992 | 546.11 |
| C25 | PromethION (Nanopore) | long | 39.33 | 8.18 | 8.5 | 3326 | 130.15 |
| C20 | NovaSeq (Illumina) | 2x150 | 79.89 | 2 x 267.27 | - | - | - |
| C20 | PromethION (Nanopore) | long | 46.87 | 9.1 | 8.3 | 3780 | 857.77 |

**Supplementary Table 3.** Archaeal marker genes that were excluded in the Lokiarchaeota specific marker-gene set.

|  |  |  |
| --- | --- | --- |
| PF04010 | Protein of unknown function (DUF357) | Missing in 3 most complete genomes |
| PF01866 | Putative diphthamide synthesis protein | Missing in 3 most complete genomes |
| TIGR00289 | TIGR00289 family protein | Missing in 3 most complete genomes |
| TIGR03679 | Universal archaeal metal-binding-domain/4Fe-4S-binding-domain containing ABC transporter, ATP-binding protein | Missing in 3 most complete genomes |
| TIGR00336 | pyrE: orotate phosphoribosyltransferase | Missing in 3 most complete genomes |
| TIGR00522 | dph5: diphthine synthase | Missing in 3 most complete genomes |
| TIGR03677 | rpl7ae: 50S ribosomal protein L7Ae | Missing in 3 most complete genomes |
| TIGR03685 | L12P_arch: 50S ribosomal protein L12P | Missing in 3 most complete genomes |
| PF01287 | Eukaryotic elongation factor 5A hypusine, DNA-binding OB fold | Missing in 3 most complete genomes |
| PF00679 | Elongation factor G C-terminus | Duplicated in 3 most complete genomes |
| PF00867 | XPG I-region | Duplicated in 3 most complete genomes |
| PF00958 | GMP synthase C terminal domain | Duplicated in 3 most complete genomes |

**Supplementary Table 4.** Eukaryotic signature proteins identified in the L04 hybrid genome.

| **Locus_tag** | **Category** | **Description** |
| --- | --- | --- |
| AB25LSR04mt_03280 | Cytoskeleton | Actin family |
| AB25LSR04mt_26280 | Cytoskeleton | Actin family |
| AB25LSR04mt_00850 | Cytoskeleton | Actin family |
| AB25LSR04mt_02580 | Cytoskeleton | Actin family |
| AB25LSR04mt_08630 | Cytoskeleton | Actin family |
| AB25LSR04mt_31940 | Cytoskeleton | Actin family |
| AB25LSR04mt_29800 | Cytoskeleton | Cell division GTPase (FtsZ) |
| AB25LSR04mt_28250 | Cytoskeleton | Gelsolin-like domain |
| AB25LSR04mt_29700 | Cytoskeleton | hypothetical proteins assigned to same arCOG like Profilin-domain proteins |
| AB25LSR04mt_00920 | Cytoskeleton | Profilin domain |
| AB25LSR04mt_05740 | Cytoskeleton | Profilin domain |
| AB25LSR04mt_31730 | Cytoskeleton | Profilin domain |
| AB25LSR04mt_34100 | Cytoskeleton | Profilin domain |
| AB25LSR04mt_04700 | Cytoskeleton | putative cytoskeleton/cell division |
| AB25LSR04mt_09460 | Cytoskeleton | putative cytoskeleton/cell division |
| AB25LSR04mt_15670 | Cytoskeleton | putative cytoskeleton/cell division |
| AB25LSR04mt_02630 | Cytoskeleton | Villin/Gelsolin |
| AB25LSR04mt_14650 | Cytoskeleton | Villin/Gelsolin |
| AB25LSR04mt_21020 | Cytoskeleton | Villin/Gelsolin |
| AB25LSR04mt_27050 | Cytoskeleton | Villin/Gelsolin |
| AB25LSR04mt_37760 | Cytoskeleton | Villin/Gelsolin |
| AB25LSR04mt_16300 | ESCRT | EAP30 domain protein (ESCRT-II) (Vps22/36-like) |
| AB25LSR04mt_16290 | ESCRT | ESCRT-II complex, Vps25 subunit, N-terminal winged helix; ESCRT-II complex, vps25 subunit; Winged helix-turn-helix DNA-binding domain |
| AB25LSR04mt_16270 | ESCRT | SNF7 family protein |
| AB25LSR04mt_22660 | ESCRT | SNF7 family protein |
| AB25LSR04mt_22670 | ESCRT | Steadiness box (SB) domain |
| AB25LSR04mt_27280 | ESCRT | Vacuolar protein sorting-associated, VPS28 |
| AB25LSR04mt_16280 | ESCRT | Vps4 ATPase with characteristic MIT domain |
| AB25LSR04mt_30410 | ESCRT | Vps4 ATPase with characteristic MIT domain |
| AB25LSR04mt_31400 | ESCRT | Vps4 ATPase with characteristic MIT domain |
| AB25LSR04mt_33730 | ESCRT | Vps4 ATPase with characteristic MIT domain |
| AB25LSR04mt_37540 | ESCRT | Vps4 ATPase with characteristic MIT domain |
| AB25LSR04mt_03730 | gtpases |  |
| AB25LSR04mt_00590 | gtpases | gtpases |
| AB25LSR04mt_00750 | gtpases | gtpases |
| AB25LSR04mt_01320 | gtpases | gtpases |
| AB25LSR04mt_01910 | gtpases | gtpases |
| AB25LSR04mt_02520 | gtpases | gtpases |
| AB25LSR04mt_03170 | gtpases | gtpases |
| AB25LSR04mt_04460 | gtpases | gtpases |
| AB25LSR04mt_04540 | gtpases | gtpases |
| AB25LSR04mt_05570 | gtpases | gtpases |
| AB25LSR04mt_05780 | gtpases | gtpases |
| AB25LSR04mt_07350 | gtpases | gtpases |
| AB25LSR04mt_07570 | gtpases | gtpases |
| AB25LSR04mt_08430 | gtpases | gtpases |
| AB25LSR04mt_09140 | gtpases | gtpases |
| AB25LSR04mt_09270 | gtpases | gtpases |
| AB25LSR04mt_09310 | gtpases | gtpases |
| AB25LSR04mt_09350 | gtpases | gtpases |
| AB25LSR04mt_09440 | gtpases | gtpases |
| AB25LSR04mt_10140 | gtpases | gtpases |
| AB25LSR04mt_10240 | gtpases | gtpases |
| AB25LSR04mt_11640 | gtpases | gtpases |
| AB25LSR04mt_12070 | gtpases | gtpases |
| AB25LSR04mt_13980 | gtpases | gtpases |
| AB25LSR04mt_14000 | gtpases | gtpases |
| AB25LSR04mt_14050 | gtpases | gtpases |
| AB25LSR04mt_14610 | gtpases | gtpases |
| AB25LSR04mt_16120 | gtpases | gtpases |
| AB25LSR04mt_17680 | gtpases | gtpases |
| AB25LSR04mt_17770 | gtpases | gtpases |
| AB25LSR04mt_17990 | gtpases | gtpases |
| AB25LSR04mt_18410 | gtpases | gtpases |
| AB25LSR04mt_18420 | gtpases | gtpases |
| AB25LSR04mt_18430 | gtpases | gtpases |
| AB25LSR04mt_18560 | gtpases | gtpases |
| AB25LSR04mt_18820 | gtpases | gtpases |
| AB25LSR04mt_19050 | gtpases | gtpases |
| AB25LSR04mt_19180 | gtpases | gtpases |
| AB25LSR04mt_20010 | gtpases | gtpases |
| AB25LSR04mt_20030 | gtpases | gtpases |
| AB25LSR04mt_21160 | gtpases | gtpases |
| AB25LSR04mt_21470 | gtpases | gtpases |
| AB25LSR04mt_21640 | gtpases | gtpases |
| AB25LSR04mt_21800 | gtpases | gtpases |
| AB25LSR04mt_21920 | gtpases | gtpases |
| AB25LSR04mt_21950 | gtpases | gtpases |
| AB25LSR04mt_22170 | gtpases | gtpases |
| AB25LSR04mt_22420 | gtpases | gtpases |
| AB25LSR04mt_22510 | gtpases | gtpases |
| AB25LSR04mt_22730 | gtpases | gtpases |
| AB25LSR04mt_22940 | gtpases | gtpases |
| AB25LSR04mt_23490 | gtpases | gtpases |
| AB25LSR04mt_23990 | gtpases | gtpases |
| AB25LSR04mt_26010 | gtpases | gtpases |
| AB25LSR04mt_26200 | gtpases | gtpases |
| AB25LSR04mt_28450 | gtpases | gtpases |
| AB25LSR04mt_29330 | gtpases | gtpases |
| AB25LSR04mt_29350 | gtpases | gtpases |
| AB25LSR04mt_29540 | gtpases | gtpases |
| AB25LSR04mt_30030 | gtpases | gtpases |
| AB25LSR04mt_30340 | gtpases | gtpases |
| AB25LSR04mt_30420 | gtpases | gtpases |
| AB25LSR04mt_30440 | gtpases | gtpases |
| AB25LSR04mt_30670 | gtpases | gtpases |
| AB25LSR04mt_30880 | gtpases | gtpases |
| AB25LSR04mt_31880 | gtpases | gtpases |
| AB25LSR04mt_32140 | gtpases | gtpases |
| AB25LSR04mt_32240 | gtpases | gtpases |
| AB25LSR04mt_32580 | gtpases | gtpases |
| AB25LSR04mt_32600 | gtpases | gtpases |
| AB25LSR04mt_32660 | gtpases | gtpases |
| AB25LSR04mt_32870 | gtpases | gtpases |
| AB25LSR04mt_33350 | gtpases | gtpases |
| AB25LSR04mt_33700 | gtpases | gtpases |
| AB25LSR04mt_33790 | gtpases | gtpases |
| AB25LSR04mt_33840 | gtpases | gtpases |
| AB25LSR04mt_34010 | gtpases | gtpases |
| AB25LSR04mt_34450 | gtpases | gtpases |
| AB25LSR04mt_34890 | gtpases | gtpases |
| AB25LSR04mt_35900 | gtpases | gtpases |
| AB25LSR04mt_36160 | gtpases | gtpases |
| AB25LSR04mt_37040 | gtpases | gtpases |
| AB25LSR04mt_37370 | gtpases | gtpases |
| AB25LSR04mt_37420 | gtpases | gtpases |
| AB25LSR04mt_37840 | gtpases | gtpases |
| AB25LSR04mt_38130 | gtpases | gtpases |
| AB25LSR04mt_38260 | gtpases | gtpases |
| AB25LSR04mt_27790 | OST | Oligosaccharyl transferase complex, subunit OST3/OST6 |
| AB25LSR04mt_19980 | OST | Oligosaccharyl transferase, STT3 subunit |
| AB25LSR04mt_22070 | OST | Oligosaccharyl transferase, STT3 subunit |
| AB25LSR04mt_28090 | OST | Ribophorin I |
| AB25LSR04mt_29440 | Trafficking machinery |  |
| AB25LSR04mt_15680 | Trafficking machinery | Coatomer, epsilon subunit |
| AB25LSR04mt_18000 | Trafficking machinery | Coatomer, epsilon subunit |
| AB25LSR04mt_21500 | Trafficking machinery | Coatomer, epsilon subunit |
| AB25LSR04mt_08030 | Trafficking machinery | Longin-like domain and MON1 domain |
| AB25LSR04mt_16040 | Trafficking machinery | Longin-like domain and MON1 domain |
| AB25LSR04mt_18610 | Trafficking machinery | Longin-like domain and MON1 domain |
| AB25LSR04mt_21130 | Trafficking machinery | Longin-like domain and MON1 domain |
| AB25LSR04mt_33290 | Trafficking machinery | Longin-like domain and MON1 domain |
| AB25LSR04mt_02310 | Trafficking machinery | Roadblock/LAMTOR2 domain; (Dynein light chain-related) (roadblock/LC7 domain) |
| AB25LSR04mt_02620 | Trafficking machinery | Roadblock/LAMTOR2 domain; (Dynein light chain-related) (roadblock/LC7 domain) |
| AB25LSR04mt_02930 | Trafficking machinery | Roadblock/LAMTOR2 domain; (Dynein light chain-related) (roadblock/LC7 domain) |
| AB25LSR04mt_04320 | Trafficking machinery | Roadblock/LAMTOR2 domain; (Dynein light chain-related) (roadblock/LC7 domain) |
| AB25LSR04mt_06840 | Trafficking machinery | Roadblock/LAMTOR2 domain; (Dynein light chain-related) (roadblock/LC7 domain) |
| AB25LSR04mt_06850 | Trafficking machinery | Roadblock/LAMTOR2 domain; (Dynein light chain-related) (roadblock/LC7 domain) |
| AB25LSR04mt_10190 | Trafficking machinery | Roadblock/LAMTOR2 domain; (Dynein light chain-related) (roadblock/LC7 domain) |
| AB25LSR04mt_21610 | Trafficking machinery | Roadblock/LAMTOR2 domain; (Dynein light chain-related) (roadblock/LC7 domain) |
| AB25LSR04mt_21670 | Trafficking machinery | Roadblock/LAMTOR2 domain; (Dynein light chain-related) (roadblock/LC7 domain) |
| AB25LSR04mt_22750 | Trafficking machinery | Roadblock/LAMTOR2 domain; (Dynein light chain-related) (roadblock/LC7 domain) |
| AB25LSR04mt_25490 | Trafficking machinery | Roadblock/LAMTOR2 domain; (Dynein light chain-related) (roadblock/LC7 domain) |
| AB25LSR04mt_27460 | Trafficking machinery | Roadblock/LAMTOR2 domain; (Dynein light chain-related) (roadblock/LC7 domain) |
| AB25LSR04mt_28180 | Trafficking machinery | Roadblock/LAMTOR2 domain; (Dynein light chain-related) (roadblock/LC7 domain) |
| AB25LSR04mt_28800 | Trafficking machinery | Roadblock/LAMTOR2 domain; (Dynein light chain-related) (roadblock/LC7 domain) |
| AB25LSR04mt_30020 | Trafficking machinery | Roadblock/LAMTOR2 domain; (Dynein light chain-related) (roadblock/LC7 domain) |
| AB25LSR04mt_31210 | Trafficking machinery | Roadblock/LAMTOR2 domain; (Dynein light chain-related) (roadblock/LC7 domain) |
| AB25LSR04mt_02030 | Ubiquitin | E3 UFM1-protein ligase 1 |
| AB25LSR04mt_00700 | Ubiquitin | JAB1/MPN/MOV34 metalloenzyme domain |
| AB25LSR04mt_04250 | Ubiquitin | JAB1/MPN/MOV34 metalloenzyme domain |
| AB25LSR04mt_04260 | Ubiquitin | JAB1/MPN/MOV34 metalloenzyme domain |
| AB25LSR04mt_04290 | Ubiquitin | JAB1/MPN/MOV34 metalloenzyme domain |
| AB25LSR04mt_26440 | Ubiquitin | JAB1/MPN/MOV34 metalloenzyme domain |
| AB25LSR04mt_34310 | Ubiquitin | JAB1/MPN/MOV34 metalloenzyme domain |
| AB25LSR04mt_01290 | Ubiquitin | THIF-type NAD/FAD binding fold; Ubiquitin-activating enzyme, catalytic cysteine domain; E2 binding; Ubiquitin activating enzyme, alpha domain; |
| AB25LSR04mt_20070 | Ubiquitin | THIF-type NAD/FAD binding fold; Ubiquitin-activating enzyme, catalytic cysteine domain; E2 binding; Ubiquitin activating enzyme, alpha domain; |
| AB25LSR04mt_26420 | Ubiquitin | THIF-type NAD/FAD binding fold; Ubiquitin-activating enzyme, catalytic cysteine domain; E2 binding; Ubiquitin activating enzyme, alpha domain; |
| AB25LSR04mt_00010 | Ubiquitin | Ubiquitin-conjugating enzyme E2; Ubiquitin-conjugating enzyme/RWD-like; Ubiquitin-conjugating enzyme, active site |
| AB25LSR04mt_20100 | Ubiquitin | Ubiquitin-conjugating enzyme E2; Ubiquitin-conjugating enzyme/RWD-like; Ubiquitin-conjugating enzyme, active site |
| AB25LSR04mt_26430 | Ubiquitin | Ubiquitin-conjugating enzyme E2; Ubiquitin-conjugating enzyme/RWD-like; Ubiquitin-conjugating enzyme, active site |
| AB25LSR04mt_07210 | Ubiquitin | Ubiquitin-related domain; Ubiquitin-like; Ubiquitin |
| AB25LSR04mt_26350 | Ubiquitin | Ubiquitin-related domain; Ubiquitin-like; Ubiquitin |
| AB25LSR04mt_26380 | Ubiquitin | Ubiquitin-related domain; Ubiquitin-like; Ubiquitin |
| AB25LSR04mt_26410 | Ubiquitin | Ubiquitin-related domain; Ubiquitin-like; Ubiquitin |
| AB25LSR04mt_04760 | Ubiquitin | Zinc finger, RING/FYVE/PHD-type |
| AB25LSR04mt_06880 | Ubiquitin | Zinc finger, RING/FYVE/PHD-type |
| AB25LSR04mt_11520 | Ubiquitin | Zinc finger, RING/FYVE/PHD-type |
| AB25LSR04mt_16850 | Ubiquitin | Zinc finger, RING/FYVE/PHD-type |
| AB25LSR04mt_20960 | Ubiquitin | Zinc finger, RING/FYVE/PHD-type |
| AB25LSR04mt_26910 | Ubiquitin | Zinc finger, RING/FYVE/PHD-type |
| AB25LSR04mt_27770 | Ubiquitin | Zinc finger, RING/FYVE/PHD-type |
| AB25LSR04mt_29590 | Ubiquitin | Zinc finger, RING/FYVE/PHD-type |
| AB25LSR04mt_33660 | Ubiquitin | Zinc finger, RING/FYVE/PHD-type |
| AB25LSR04mt_37430 | Ubiquitin | Zinc finger, RING/FYVE/PHD-type |

**Supplementary Table 5.** Number of ESPs identified in the L04 hybrid genome per category.

|  | *Lokiarchaeum*<br>GC14_75 | Lokiarchaeote_CR_<br>4 | Hybrid L04 | MK-D1 |
| --- | --- | --- | --- | --- |
| **Cytoskeleton** | 23 | 13 | 21 | 15 |
| **Ubiquitin** | 32 | 26 | 27 | 15 |
| **ESCRT** | 9 | 8 | 11 | 11 |
| **Trafficking**<br>**machinery** | 26 | 23 | 25 | 21 |
| **GTPases** | 117 | 84 | 87 | 64 |
| **OST** | 4 | 3 | 4 | 4 |
|  | 211 | 157 | 185 | 131 |

**Supplementary Table 6:** Genomic regions including bases no covered by any read. If several bases in close proximity had 0 read coverage, a window spanning neighbouring 0-coverage bases is reported.

| Contig_id | From | To | Length of the window |
| --- | --- | --- | --- |
| **Regions including bases no covered by any read using the unfiltered long-read alignments** |  |  |  |
| loki04_scf7180000000547_pilon_pilon_pilon | 2140485 | 2140585 | 100 |
| loki04_scf7180000000493_pilon_pilon_pilon | 188929 | 190907 | 1978 |
| **Regions including bases no covered by any read using the filtered long-read alignments** |  |  |  |
| loki04_scf7180000000547_pilon_pilon_pilon | 35 | 10989 | 10954 |
| loki04_scf7180000000547_pilon_pilon_pilon | 153748 | 153749 | 1 |
| loki04_scf7180000000547_pilon_pilon_pilon | 1514258 | 1521754 | 7496 |
| loki04_scf7180000000547_pilon_pilon_pilon | 2137687 | 2146130 | 8443 |
| loki04_scf7180000000484_pilon_pilon_pilon | 0 | 921 | 921 |
| loki04_scf7180000000484_pilon_pilon_pilon | 583094 | 583533 | 439 |
| loki04_scf7180000000484_pilon_pilon_pilon | 1052996 | 1053546 | 550 |
| loki04_scf7180000000484_pilon_pilon_pilon | 1065225 | 1065226 | 1 |
| loki04_scf7180000000484_pilon_pilon_pilon | 1071778 | 1076594 | 4816 |
| loki04_scf7180000000484_pilon_pilon_pilon | 1723461 | 1729042 | 5581 |
| loki04_scf7180000000493_pilon_pilon_pilon | 0 | 9087 | 9087 |
| loki04_scf7180000000493_pilon_pilon_pilon | 188916 | 192821 | 3905 |
| **Regions including bases no covered by any read using the short-read alignments** |  |  |  |
| loki04_scf7180000000547_pilon_pilon_pilon | 5258 | 15647 | 10389 |
| loki04_scf7180000000547_pilon_pilon_pilon | 56004 | 56038 | 34 |
| loki04_scf7180000000547_pilon_pilon_pilon | 234419 | 234420 | 1 |
| loki04_scf7180000000547_pilon_pilon_pilon | 295211 | 295235 | 24 |
| loki04_scf7180000000547_pilon_pilon_pilon | 914793 | 914794 | 1 |
| loki04_scf7180000000547_pilon_pilon_pilon | 1386335 | 1386342 | 7 |
| loki04_scf7180000000547_pilon_pilon_pilon | 1518852 | 1518853 | 1 |
| loki04_scf7180000000547_pilon_pilon_pilon | 1661914 | 1661915 | 1 |
| loki04_scf7180000000547_pilon_pilon_pilon | 1703218 | 1703227 | 9 |
| loki04_scf7180000000547_pilon_pilon_pilon | 1810439 | 1810455 | 16 |
| loki04_scf7180000000547_pilon_pilon_pilon | 2140485 | 2143343 | 2858 |
| loki04_scf7180000000484_pilon_pilon_pilon | 0 | 51 | 51 |
| loki04_scf7180000000484_pilon_pilon_pilon | 544921 | 544951 | 30 |
| loki04_scf7180000000484_pilon_pilon_pilon | 566002 | 566013 | 11 |
| loki04_scf7180000000484_pilon_pilon_pilon | 848803 | 848842 | 39 |
| loki04_scf7180000000484_pilon_pilon_pilon | 913913 | 917300 | 3387 |
| loki04_scf7180000000484_pilon_pilon_pilon | 1054610 | 1054611 | 1 |
| loki04_scf7180000000484_pilon_pilon_pilon | 1063703 | 1063809 | 106 |
| loki04_scf7180000000484_pilon_pilon_pilon | 1076444 | 1076559 | 115 |
| loki04_scf7180000000484_pilon_pilon_pilon | 1373340 | 1373356 | 16 |
| loki04_scf7180000000484_pilon_pilon_pilon | 1506034 | 1506056 | 22 |
| loki04_scf7180000000493_pilon_pilon_pilon | 4345 | 6708 | 2363 |
| loki04_scf7180000000493_pilon_pilon_pilon | 172958 | 172968 | 10 |
| loki04_scf7180000000493_pilon_pilon_pilon | 188929 | 191041 | 2112 |

